# Single-trial neural dynamics influence auditory category learning

**DOI:** 10.1101/2020.12.10.420091

**Authors:** Kelsey Mankel, Philip I. Pavlik, Gavin M. Bidelman

## Abstract

Percepts are naturally grouped into meaningful categories to process continuous stimulus variations in the environment. Theories of category acquisition have existed for decades, but how they arise in the brain due to learning is not well understood. Here, advanced computational modeling techniques borrowed from educational data mining and cognitive psychology were used to trace the development of auditory categories within a short-term training session. Nonmusicians were rapidly trained for 20 min on musical interval identification (i.e., minor and major 3^rd^ interval dyads) while their brain activity was recorded via EEG. Categorization performance and neural responses were then assessed for the trained (3^rds^) and novel untrained (major/minor 6^ths^) continua. Computational modeling was used to predict behavioral identification responses and whether the inclusion of single-trial features of the neural data could predict successful learning performance. Model results revealed meaningful brain-behavior relationships in auditory category learning detectible on the single-trial level; smaller P2 amplitudes were associated with a greater probability of correct interval categorization after learning. These findings highlight the nuanced dynamics of brain-behavior coupling that help explain the temporal emergence of auditory categorical learning in the brain.

## I. INTRODUCTION

To make sense of the environment, percepts are naturally grouped into meaningful categories, a phenomenon known as categorical perception (CP; Harnad, 1987). Category acquisition is generally believed to involve both innate and learned components (Rosen and Howell, 1987). While long-term plasticity in auditory categorization is well documented (e.g., musical training: Zatorre and Halpern, 1979; language experience: Kuhl, 1991; Kuhl *et al.*, 1992), how categories develop through *short-term* learning is less understood.

Behaviorally, musical interval identification improves with training (Pavlik Jr *et al.*, 2013; Little *et al.*, 2019) as do behavioral thresholds after pitch discrimination training (Carcagno and Plack, 2011). Even nonmusicians can improve in musical interval identification (Pavlik Jr *et al.*, 2013; Little *et al.*, 2019) and discrimination (Burns and Ward, 1978), suggesting that learning, rather than music experience *per se*, promotes the successful labeling of musical sounds. Similarly, short-term training on non-native phonetic contrasts leads to behavioral and neural improvements in speech identification and discrimination after learning (Pisoni *et al.*, 1982; Lively *et al.*, 1993; Kraus *et al.*, 1995; Tremblay *et al.*, 2001; Myers and Swan, 2012; Swan and Myers, 2013). Category learning is thought to distort the stimulus representational space, such that auditory cortical maps emphasize differences between categories and become more insensitive to within-category differences (Guenther *et al.*, 1999; Guenther *et al.*, 2004; e.g., see Fig. 1 in Bidelman *et al.*, 2020).

**Fig. 1:**
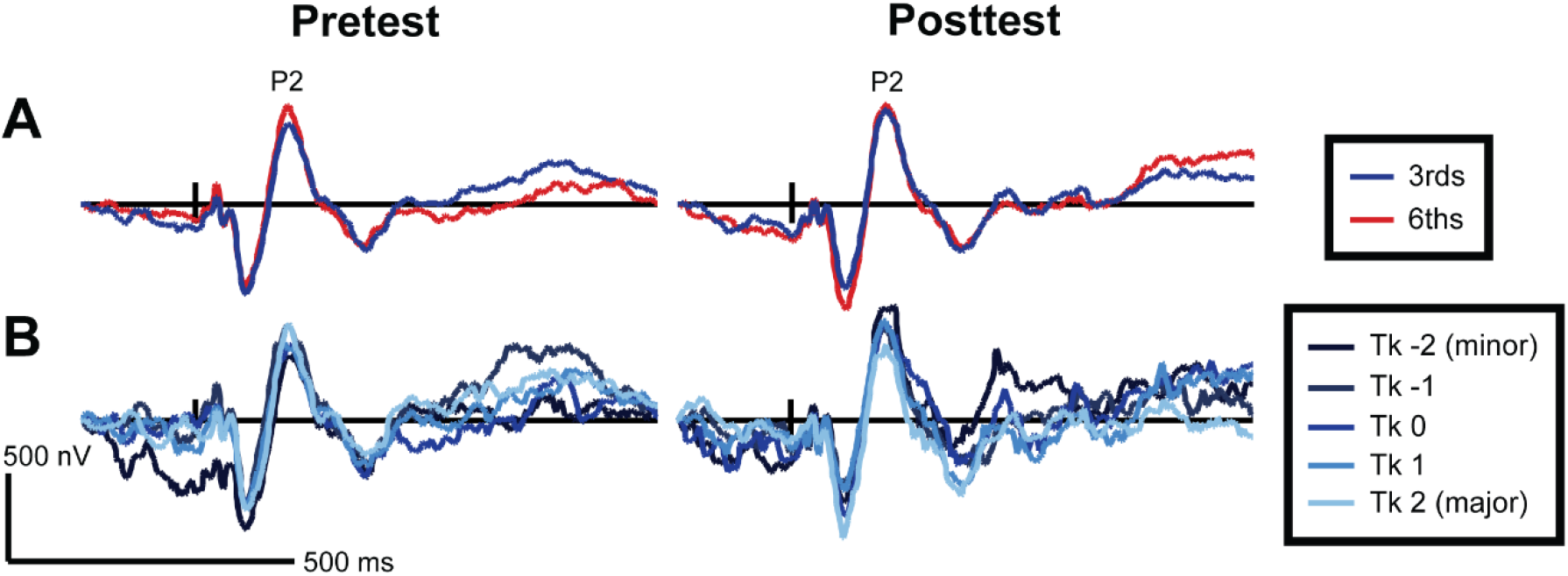
(A) Grand average ERP responses for pre- and posttest, separated by 3^rd^ vs. 6^th^ intervals, reflecting averaged activity across all 5 tokens per interval continuum (*n*=20). (B) Grand average responses for the 3^rds^ interval continuum indicate categorization differences after training (posttest) around the timeframe of P2 (~150-200 ms), particularly for the minor vs. major prototypical tokens (i.e., tokens −2 and 2, respectively). Color figures are available in the online version of this manuscript.

Event-related potentials (ERPs) have been instrumental in shedding light on the neural mechanisms and time course of categorization processes. M/EEG studies have demonstrated that speech categories begin to emerge around N1 (~100 ms post-stimulus onset) and are fully formed by P2 (~ 150-200 ms) (Bidelman *et al.*, 2013b; Ross *et al.*, 2013; Bidelman *et al.*, 2014; Bidelman and Alain, 2015; Alho *et al.*, 2016; Bidelman and Walker, 2017; 2019; Mankel *et al.*, 2020). Source localization studies have invoked a distributed frontotemporal network involved in speech categorization, including key brain regions such as the left primary auditory cortex (PAC), superior temporal gyrus (STG), and inferior frontal gyrus (IFG) (Binder *et al.*, 2004; Golestani and Zatorre, 2004; Liebenthal *et al.*, 2005; Desai *et al.*, 2008; Myers *et al.*, 2009; Chang *et al.*, 2010; Liebenthal *et al.*, 2010; Lee *et al.*, 2012; Myers and Swan, 2012; Alho *et al.*, 2016; Bouton *et al.*, 2018; Bidelman and Walker, 2019; Mankel *et al.*, 2020). The neural underpinnings of music categorization have been less studied, but results suggest a similar (albeit right hemisphere biased) network involving the right STG/STS (Klein and Zatorre, 2011; 2015; Bidelman and Walker, 2019; Mankel *et al.*, 2020). Several studies have also reported experience-dependent changes in CP at behavioral and neural levels associated with music training (Bidelman *et al.*, 2014; Bidelman and Alain, 2015; Wu *et al.*, 2015; Bidelman and Walker, 2017; 2019) and tone language expertise (Bidelman and Lee, 2015), underscoring the role of *long-term* auditory experience in categorization processes.

*Short-term* learning-related changes in nonnative phonetic perception have been associated with changes in P2 (and magnetic P2m) as well as late slow activity (~250-400 ms) of the ERPs (Tremblay *et al.*, 2001; Alain *et al.*, 2010; Ben-David *et al.*, 2011; Carcagno and Plack, 2011; Ross *et al.*, 2013). Some report decreased or more efficient P2 responses after training (Golestani and Zatorre, 2004; Alain *et al.*, 2010; Ben-David *et al.*, 2011) while others show robust increases in amplitudes (Draganova *et al.*, 2009; Tong *et al.*, 2009; Ross *et al.*, 2013). Rapid changes in both temporal (e.g., STG) and frontal (e.g., IFG) brain areas have also been observed following short-term auditory discrimination training (de Souza *et al.*, 2013), nonspeech categorization training (Guenther *et al.*, 2004), task-related improvements in concurrent speech segregation (Alain *et al.*, 2007; Du *et al.*, 2015), and tone language learning (Lee *et al.*, 2017). Collectively, the extant literature indicates that successful short-term auditory category learning is accompanied by neurophysiological changes around the time frame of P2 (i.e., within 150-250 ms).

Much of the existing neuroscience and psychological research on auditory categorization have focused on characterizing averaged outcomes rather than modeling or making predictions about single event data (Yarkoni and Westfall, 2017). While averaging trial data reduces noise, it may wash out underlying patterns in the data, particularly those related to individual differences and the temporal dynamics of performance. Computational modeling has been used in various areas of cognitive psychology, educational data mining, and intelligent tutoring systems to trace knowledge development, model underlying learning behavior mathematically, and predict and optimize human learning performance (Pavlik Jr. and Anderson, 2008; Khajah *et al.*, 2014). The generalized knowledge tracing (GKT) framework assumes that learning can be quantified in terms of knowledge components that depict learning difficulty for a particular item or skill via logistic regression models, typically with binary outcomes such as accuracy on a particular test item (Spada and McGaw, 1985; Pavlik Jr *et al.*, 2020). The GKT framework allows implementation of models such as the additive factors model (AFM; Cen *et al.*, 2006) and performance factors analysis model (PFA; Pavlik Jr *et al.*, 2009). Both of these learning models factor the quantity of knowledge or prior training with a knowledge component by scaling the effect of the number of prior practice trials for a particular item as an individual parameter (AFM) or fitting separate parameters for success and failure of prior practices (PFA). Nonlinear features, such as recency-weighted successes and/or failures (Gong *et al.*, 2011; Galyardt and Goldin, 2015) or the natural log of the successful to failure trials ratio (Pavlik Jr *et al.*, 2020), might also add further explanatory power to GKT models. Because such models predict single responses in a task, they are capable of representing an individual’s knowledge acquisition and learning rate, provided there is enough data for reliable parameter estimates (Liu and Koedinger, 2017; Pavlik Jr *et al.*, 2020).

In this study, we were interested in characterizing the short-term neuroplasticity that manifests during rapid auditory category learning. Adopting aspects of the GKT framework for a novel view into category acquisition, we modeled rapid auditory category learning to trace the development of musical interval categories at brain and behavioral levels. Listeners were trained to identify musical intervals as they are not overlearned (cf. speech) and thus represent relatively novel stimuli that do not carry categorical labels for nonmusicians (Burns and Ward, 1978; Zatorre and Halpern, 1979; Bidelman and Walker, 2017). Parameters were included in the computational models based on listeners’ single-trial EEG responses to assess whether inclusion of neural data improves model predictions. We hypothesized that (1) category learning would develop rapidly within a short (20 min) training session; (2) learning would transfer to untrained stimuli (i.e., music intervals not present in the learning phase); and (3) incorporating neural measures in the models (i.e., ERP P2) would yield better predictions of learning outcomes than models based on behavior alone.

## II. METHODS

### A. Participants

Twenty young adults (*μ*=25.2 ± 4.0 yrs, 16 females) were recruited for this study. All participants had normal hearing (<25 dB SPL, 250-8000 Hz), were right-handed (Oldfield, 1971), and had no history of neurological disorders. Participants were required to be fluent speakers of English; 6 reported a native language other than English according to language history questionnaires (Li *et al.*, 2006). Importantly, none of the participants had any tone language experience as these languages improve musical pitch perception (Bidelman *et al.*, 2013a). All participants had minimal to no formal music experience (μ=1.1 ± 1.1 yrs, <3 years on any combination of instruments) and were thus naïve to the music-theoretic labels for pitch intervals. Participants gave written informed consent according to protocol approved by the University of Memphis Institutional Review Board, and they were compensated $10 per hour for their time (~2.5-3 hours total duration).

### B. Stimuli

Two five-step musical interval continua were constructed of complex tones consisting of 10 equal amplitude harmonics added in cosine phase. The fundamental frequency for the bass note across both continua was fixed at 150 Hz, while the upper note of the harmonic interval ranged from 180-188 Hz (spanning a minor to major 3^rd^) or 240-250 Hz (minor to major 6^th^) with equidistant frequency spacing between adjacent steps along the continuum. These intervals were selected because both continua span a semitone and are considered similar in qualia (i.e., imperfect consonances) in Western music practice. Each token was 100 ms in duration with a 10 ms rise/fall time to reduce spectral splatter. Stimulus presentation was controlled via MATLAB and routed through a TDT interface (Tucker Davis Technologies).

### C. Procedure

Subjects were seated comfortably in an electroacoustically shielded booth. Stimuli were presented binaurally through ER-2 insert earphones at 80 dB SPL (Etymotic Research). Baseline categorization was assessed in the pretest, followed by a brief training session on the minor/major interval categories, then a posttest which measured learning-associated changes in performance. During the pre- and post-test phases, 3^rds^ and 6^ths^ were presented in separate blocks (i.e., one block for each continuum per phase, counterbalanced across participants). Approximately 2-3 minor/major exemplars were played at the beginning of each block to orient participants to the stimulus categories. For both the pretest and posttest phases, each token of the continuum was randomly presented 120 times for a total of 600 trials per block (5 tokens x 120 = 600 trials; 1200 total trials in pretest and posttest)^1^. On each trial, participants were asked to label the sounds they heard as either “minor” or “major” via keyboard button press as fast and accurately as possible. Feedback was not provided. The interstimulus interval was jittered randomly between 400-600 ms (20 ms steps, uniform distribution) following the listener’s response to avoid anticipation of the next trial, reduce rhythmic entrainment of EEG oscillations, and to help filter out overlapping activity from the previous trial (Luck, 2014). Participants were offered a break between blocks to reduce fatigue.

Between the pre- and post-test, participants performed a single, approximately 20-minute identification training session. Training consisted of 250 presentations of each exemplar (endpoint) from the minor and major 3^rd^ continuum (total = 500 trials spread across 10 blocks, 25 randomized trials of each token per block)^2^. The 6^ths^ continuum was withheld from training. Feedback was given during the training to improve accuracy and efficiency of auditory category learning (Yi and Chandrasekaran, 2016). Training only on the endpoints of the 3^rd^ continuum allowed us to examine (i) the perceptual warping of the remaining stimulus space (cf. Livingston *et al.*, 1998; Guenther *et al.*, 1999) and (ii) evaluate transfer effects to the untrained 6^th^ intervals. EEGs were recorded continuously throughout the experiment (i.e., pretest, training, & posttest phases).

### D. EEG acquisition and preprocessing

EEG data were recorded from 64 sintered Ag/AgCl electrodes at 10-10 scalp locations (Oostenveld and Praamstra, 2001) and digitized at a sampling rate of 500 Hz (Synamps RT amplifier; Compumedics Neuroscan). Electrodes were referenced during acquisition to an additional sensor placed approximately 1 cm posterior to Cz. Impedances were set to <10 kΩ at the start of data collection, and caps were refreshed with saline as needed prior to the posttest. Ocular movements were monitored by electrodes placed on the outer canthi of the eyes and the superior and inferior orbit. The data were epoched (−200-800 ms), filtered (1-30 Hz, 4^th^-order Butterworth filter), and re-referenced offline to the common average reference.

The neural correlates of auditory categorization emerge between the N1 and P2 deflections (100-150ms) (Bidelman *et al.*, 2013b; Bidelman and Alain, 2015; Bidelman and Lee, 2015; Alho *et al.*, 2016; Bidelman and Walker, 2017; Toscano *et al.*, 2018; Bidelman and Walker, 2019; Mankel *et al.*, 2020), so these ERPs were explored as possible neural model predictors of categorical learning. Single-trial ERP amplitudes and latencies were calculated as the peak negative voltage between 85-160 ms for N1 and peak positive voltage between 150-220 ms for P2 from channel Cz (Hall, 1992). N1-P2 amplitudes were computed as the difference between the individual peak amplitudes.

### E. Statistical Models

Four logistic regression models were tested in the analyses. The basic model structures are given in **Table I**, where the outcome *Response* is a logit value depicting the trial-to-trial probability of responding “major” (i.e., binary variable: 0= “minor”, 1= “major”). The predictor features for the β coefficients are written out as words to aid clarity and discussion.

**Table I.**
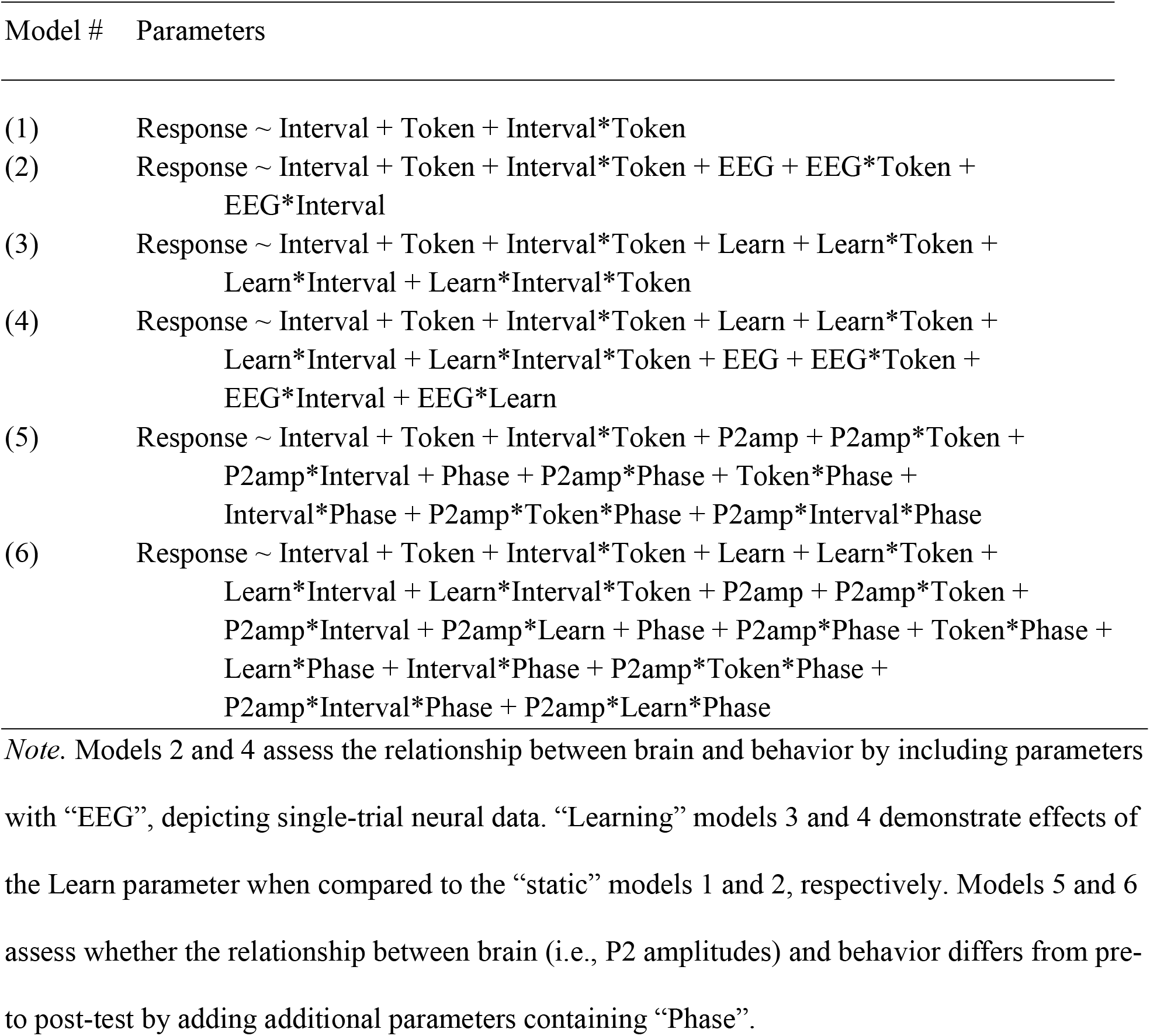
Main comparison models for predicting subject response outcomes

*Token* codes for the stimulus. These values were centered at 0 so that tokens 1-5 along the continuum ranged from −2 for minor endpoints to +2 for the major endpoints. *Interval* denotes whether the block is from the 3^rds^ or 6^ths^ continuum, which tests whether performance differs (or whether learning transfers) across stimulus sets.

Reminiscent of PFA (Pavlik Jr *et al.*, 2009), the *Learn* variable captures the accumulating history of behavioral performance as a representation of category learning. *Learn* is calculated as the natural log of the relative ratio between the summed totals of correct and incorrect trials during training, where the natural log function provides diminishing returns for learning in later trials than earlier trials (see “logit” feature: Pavlik Jr *et al.*, 2020). 1 was added to the prior trial count (i.e., both the numerator and denominator of the logit parameter) to avoid taking the log(0), which is undefined (Pavlik Jr *et al.*, 2020). *Learn* is then weighted according to the token heard on each trial throughout all phases. We assumed the prototypical musical intervals have a stronger influence over learning than more ambiguous tokens—trials with the major/minor prototypes (i.e., tokens −22, 2) are multiplied by 2, inner tokens (i.e., tokens −1, 1) are multiplied by 1, and the middle token (i.e., token 0) nulls the *Learn* variable with a multiplication factor of 0 (i.e., *Learn* = log[sum of correct trials/1 + sum of incorrect trials] x token weight). Scaling *Learn* in this way therefore allows minor and major exemplars to carry more weight in predicting responses whereas the ambiguous middle token offers equal bias on responses. Moreover, this weighting scheme captures the fact that only prototypes (and feedback) were provided in the training phase and thus contribute to categorical learning; learning gains were presumed to be constant in the pre- and post-test. Consequently, *Learn* is always 0 during pretest, increases with better accuracy during training (where only minor and major exemplars are heard), and equals the final training value multiplied by the token weight in posttest trials. Finally, *EEG* refers to the single trial EEG amplitudes or latencies for N1 and P2 (see *D. EEG acquisition and preprocessing*), where each measure was fit individually in separate model iterations. Inclusion of this neural measure assessed which ERP component most strongly contributed to behavioral learning outcomes and whether brain activity aids in predicting categorical learning on a trial-to-trial basis.

To determine the predictive performance, predictions from the model were compared to empirical data (i.e., participants’ actual responses). AIC, root mean squared error (RMSE), and McFadden’s pseudo-R^2^ values were calculated to evaluate model fit. Variance inflation factors (VIF) assessed multicollinearity between parameters during model building. Two cross-validation (CV) procedures assessed model reliability and overfitting. An “intersubject CV” involved a 10-fold, 10-run holdout method where 10% of subjects were randomly withheld for the test model while the remaining 90% were used for training (model building). A “random split CV” randomly selected 10% of the data (across all subjects) for testing in another 10-fold, 10-run holdout method. In each method, the ratio of McFadden’s R^2^ between the test and train models averaged over the 10 folds and 10 runs were computed; values closer to 1 indicate adequate reliability and minimal overfitting. Statistical analyses were completed in R (v3.5.3).

## III. RESULTS

Grand average ERPs are shown for each continuum and training phase in **Fig. 1A** while token specific responses for the 3^rds^ continuum are shown in **Fig. 1B**. We first determined which ERP components were reliable predictors of behavior. In models 2 and 4, the *EEG* parameter was replaced (separately) with single-trial N1, P2, or N1-P2 measures (see **Table I**). Models incorporating P2 latencies, N1 amplitudes and latencies, or N1-P2 amplitudes had poorer fit (i.e., smaller McFadden’s pseudo-R^2^) than those containing P2 amplitudes. Single-trial P2 amplitudes also produced larger parameter estimates compared to the other ERP components, indicating a stronger relationship between this neural measure and behavioral outcomes (Bidelman *et al.*, 2013b; Ross *et al.*, 2013; Bidelman *et al.*, 2014; Bidelman and Alain, 2015; Alho *et al.*, 2016; Bidelman and Walker, 2017; 2019; Mankel *et al.*, 2020). Thus, P2 amplitudes (*P2amp*) were used in subsequent single-trial learning models to evaluate whether brain responses inform behavioral auditory categorical learning.

Estimates, standard errors (SEs), and p-values for each of the model parameters are shown in the appendix table. Models 1 and 2 contain “static” variables that do not scale with learning. Meanwhile, models 3 and 4 contain the variable *Learn*, the log of the correct to incorrect training trials ratio (see *Methods*).

While the exact parameter estimates differ, certain patterns are evident across models. For example, the large positive estimates for *Token* demonstrate that as participants hear tokens towards the major end of the continuum (coded as +1 and +2), they are more likely to respond “major” (see Fig. 2 below). The *Interval* parameter indicates a very slight bias towards responding major for the trained 3^rds^ compared to the untrained 6^ths^. The *Interval*Token* interaction in model 2 shows a stronger effect of responding “major” for tokens towards the major end of the 3^rds^ continuum compared to the 6^ths^, suggesting that the learned categorization effects were stronger for those sounds heard during training. Meanwhile, the 3-way interaction of *Learn*Interval*Token* in model 4 demonstrates this effect was stronger for those with better training performance.

**Fig. 2:**
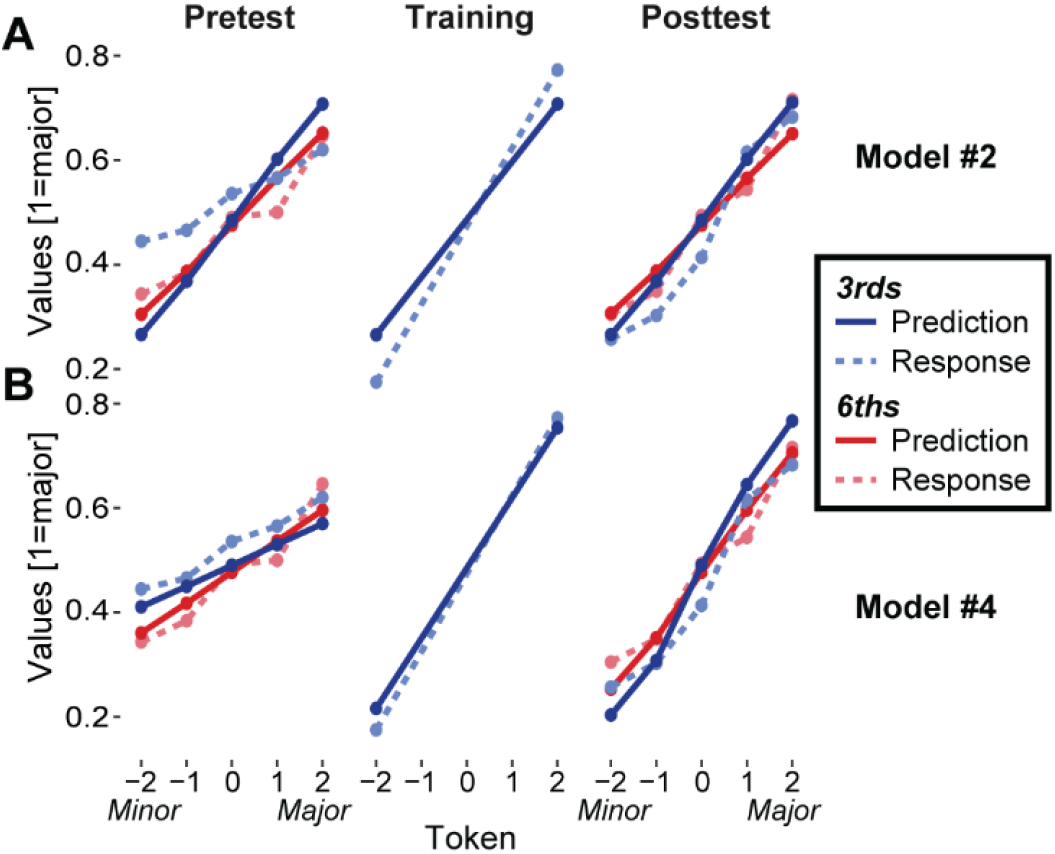
Comparisons between model predictions (solid lines) and subject behavioral responses (dashed lines) for (A) model 2 and (B) model 4 indicate better data fit for the more complex “learning” model 4 compared to the “static” model 2. Both models and behavioral responses demonstrate sharper identification curves after training, where transfer of learning is evident for the inner 3^rd^ tokens (tokens −1, 1) as well as the untrained 6^th^ continuum. Results are highly similar for the models without *P2amp* variables, models 1 and 3. Color figures are available in the online version of this manuscript.

Models 2 and 4 include the single-trial EEG data to test whether neural amplitudes (*P2amp*) are significant predictors of behavioral response outcomes. As indicated by the appendix table, *P2amp*Token* is a highly significant parameter in both the simpler static model (#2) and the more complex learning model (#4) (*p*<0.001). The *P2amp*Token* interaction reveals that larger P2 amplitudes are associated with a decreased probability of reporting “major” for tokens towards the major end of the continuum and vice versa (i.e., reduced probability of identifying minor tokens on the minor end of the continuum). The interaction of *P2amp*Interval* suggests that larger P2 amplitudes are associated with a lower probability of responding major for the trained 3^rds^ compared to the untrained 6^ths^. Additionally, *P2amp*Learn* was a significant predictor in model 4 *(p* = 0.0423), demonstrating a relationship between P2 amplitudes and successful performance.

These coefficients can be used to estimate the combined contribution of the *P2amp* variables (i.e., *P2amp, P2amp*Token, P2amp*Interval*, and *P2amp*Learn*; the latter only included in model 4) to the odds of responding major. For example, in comparing the odds of responding “major” for the major 3^rd^ token (i.e., tk 2) at an 80% accuracy rate for the training phase, a one-μV increase in P2 amplitudes corresponds to an odds ratio of 0.9468 for model 2 and 0.9669 for model 4, respectively. This means that the odds of responding “major” for the major token decreases ~4-5% with a one-μV increase in P2 amplitudes, after holding the other variables constant.

Fig. 2 compares predictions of models 2 and 4 to actual recorded responses for each token during pretest, training, and posttest. Predicted values > 0.5 are associated with higher probability of responding major, while values < 0.5 reflect minor responses. Steeper slopes of these functions indicate stronger categorization for music intervals, which is most evident in training and posttest. Training on the 3^rds^ led to stronger categorization on the inner tokens (i.e., - 1, 1). Steeper slopes from pre-to post-test are also observed for the 6^ths^, suggesting learning transferred to musical intervals not heard during training. A better correspondence between model predictions and subject responses for the dynamic (#4) rather than the static (#2) model demonstrates the effect of successful learning. Specifically, the correlation between predicted condition average values and actual average response values is larger for model 4 (*r* = 0.96) than model 2 (*r* = 0.91). Similar results were also observed for models 1 and 3, respectively.

**Table II** summarizes the fit statistics and results of the CV procedures. As expected, fits improved (i.e., larger R^2^ and smaller RMSE) for the models that included the *Learn* parameters compared to the static models. Fits also slightly improved with the addition of *P2amp.* Given that it is more difficult to predict behavior for novel subjects than to predict random trials within the same subjects, it is not surprising that the test data fits are worse for the inter-subject CV than the random split CV procedures. However, the test R^2^ and RMSEs closely approximate the respective fit measures of the full models, suggesting good predictive capabilities for new data.

The CV ratios are quite high (i.e., close to 1), indicating good correspondence between models fit to training and test data and thus rule out overfitting. VIFs were <5 for all variables in each model, suggesting limited multicollinearity among variables and more crucially, that the *Learn* and neural *P2amp* parameters captured independent variance in the data.

**Table II.**
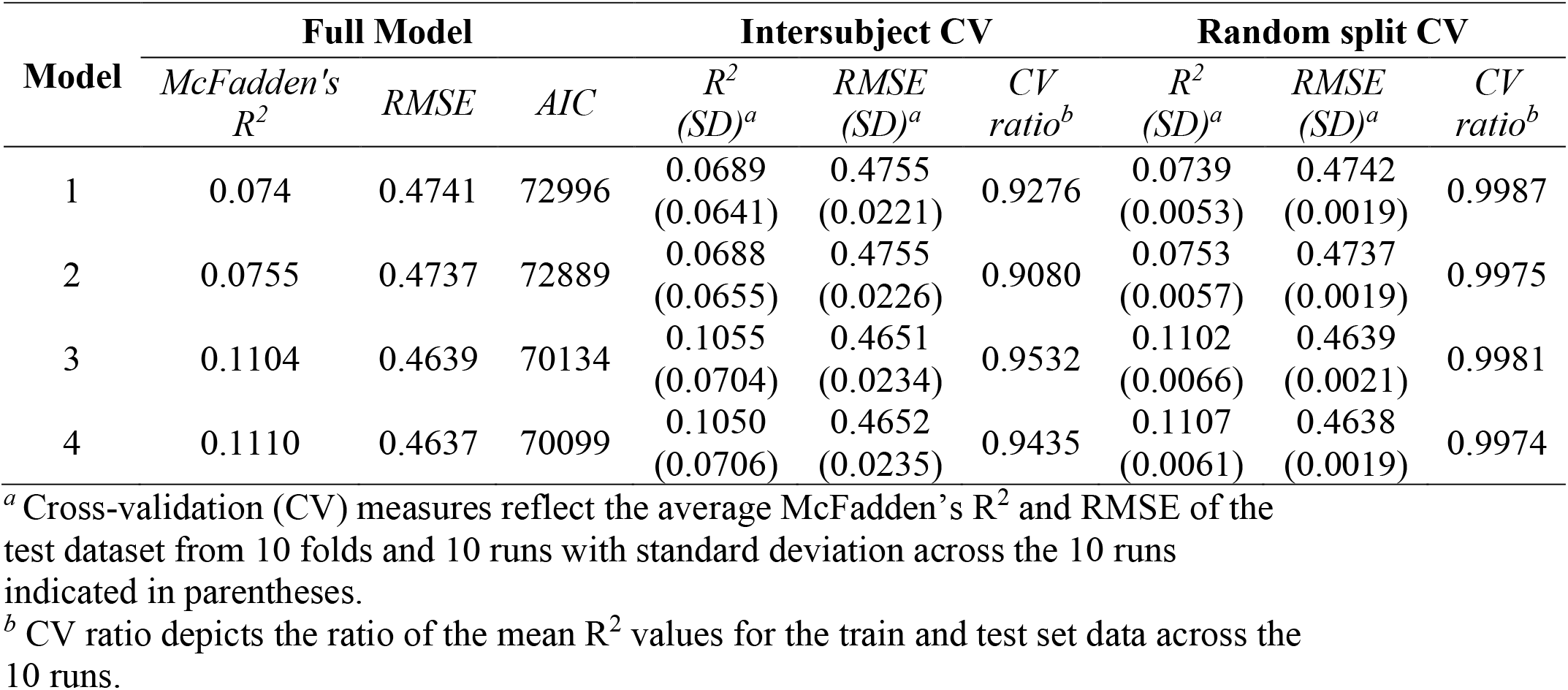
Summary of model fit and cross-validation statistics

**Fig. 3** visualizes how model predictions evolve over the course of the experiment compared to actual behavioral responses. The figure shows trial-by-trial predictions of model 4 across all three experimental phases for the major exemplar only (i.e., tk 2). The model predicts steady performance in the pre- and post-test, consistent with the general pattern of behavioral responses. In contrast, prediction values (and the proportion of “major” responses) increase steadily throughout training, where values closer to 1 correspond to a higher probability of responding “major” on each trial, indicating improvement in the ability to identify the major interval. Once these tokens are randomized among the other tokens, however, their identification is predicted to be worse in the posttest compared to the end of training, likely a result of interference from the other tokens not heard during training. Yet, performance is still better overall in the posttest compared to the pretest, demonstrating an effect of learning. Successful learning is depicted by larger values in the posttest compared to the pretest for both the 3^rds^ and, to a lesser extent, the 6^ths^, the latter effect indicating transfer. Given the binary nature of the response variable, the opposite relationship would be observed in plotting model predictions for the minor token (i.e., tk −2), whereby successful learning would be characterized by a *decrease* in values as prediction outcomes closer to 0 correspond to a higher probability of responding “minor.” These results thus demonstrate using single-trial model predictions how acquisition of a categorical knowledge component like major or minor musical intervals can be traced over time during learning.

**Fig. 3:**
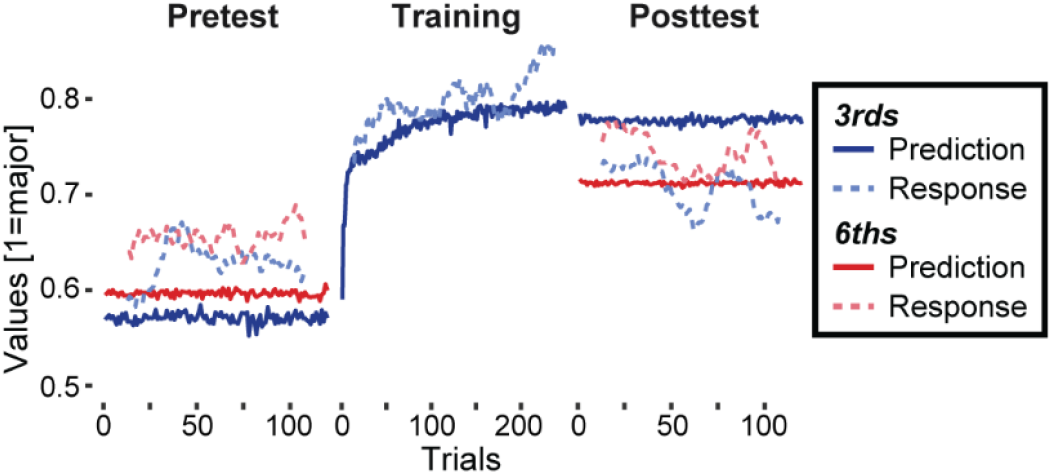
Model 4 traces the trial-to-trial development of categorical behavior throughout the experiment for the major prototypical token (tk 2). Model predictions (solid lines) and actual behavioral data (dashed lines) rise as training progresses, indicating a greater probability or proportion of responding “major” for the major token as subjects learn the appropriate interval labels. Meanwhile, the model predicts steady behavioral responses during the pre- and post-test phases, consistent with the proportion of “major” behavioral responses, but better performance is demonstrated in the posttest by higher overall prediction and response values compared to the pretest. Improvement in performance can also be seen for the untrained 6^th^ major token, though not as drastic of a pre-to post-test change in model predictions as the trained 3^rd^ token. For plotting purposes only, a running average smoother (width of 25 trials) was applied to subject responses to make the overall patterns clearer, and two subjects who terminated the 3^rds^ posttest block early due to technical issues were excluded from the figure. Color figures are available in the online version of this manuscript.

Models 2 and 4 were then fit with additional parameters to assess how the relationship between brain and behavior differs from pre-to post-test; these are depicted in Table I as models 5 and 6, respectively. We were interested in whether the interactions of *P2amp* variables with

*Phase*—dummy coded to contrast pretest vs. training or posttest trials—would suggest that the association between single-trial neural responses and behavioral outcomes changed after training. For model 5, the 3-way interaction between *Phase, P2amp*, and *Token* was significant. The negative 3-way interaction estimates suggest that the *P2amp*Token* relationship becomes stronger in both the training (β = −0.0247, *p* < 0.001) and the posttest phases (β = −0.0108, *p* = 0.0199) compared to the pretest; the effect of associating smaller P2 amplitudes with enhanced categorization (i.e., higher probability of reporting “major” for tokens on the major end of the continuum and vice versa) is enhanced after training. Both models indicated an interaction between *P2amp* and *Phase*; specifically, larger amplitudes are associated with a slightly higher probability of responding major for the training trials (β = 0.0353, *p* =0.0025), but not posttest trials (β = 0.0017, *p* = 0.8564), compared to the pretest. None of the other interactions including *P2amp* and *Phase* were significant in the more complex model 6.

## IV. DISCUSSION

We investigated whether rapid auditory category learning could be described via single-trial neural data and psychological computational models of learning. These findings show that the P2 wave of the auditory ERPs plays a significant role in predicting the gains and time-course of listeners’ perceptual learning of musical interval categories with only 20-min of training. Specifically, smaller P2 amplitudes were associated with a higher probability of correctly identifying minor and major tokens, an effect that became stronger after training. To our knowledge, this is the first study to apply learning theory models to auditory categorization and assess how the dynamics of single-trial ERP activity modulates behavioral performance during category learning.

These analyses demonstrate that even at the noisy, single-trial level, P2 amplitudes (but not other ERP components) are associated with categorization of auditory stimuli and, more critically, are linked with behavioral identification during category learning. No other ERP component was a reliable predictor of category learning in the models. This converges with other evidence suggesting that category-level representations of sound emerge by N1 and are fully formed by P2 (Bidelman *et al.*, 2013b; Ross *et al.*, 2013; Bidelman *et al.*, 2014; Bidelman and Alain, 2015; Alho *et al.*, 2016; Bidelman and Walker, 2017; 2019; Mankel *et al.*, 2020). Additionally, these results extend prior work on the neural chronometry of auditory categorization by demonstrating that meaningful brain-behavior associations develop at a single-trial neural level and are subject to rapid plasticity during short-term training.

Remarkably, category learning was evident in only one, 2.5-hour experimental session, only 20 minutes of which was spent in identification training. Specifically, smaller trial-wise P2 amplitudes were associated with a higher accuracy in identifying musical interval categories, an effect that became stronger after training. Critically, the interaction between *P2amp*Learn* further suggests this relationship is not simply exogenous (i.e., due to the brain’s mere response to stimulus properties), as the P2-behavior association scaled with successful learning. Instead, the data suggest P2 reflects more than obligatory stimulus coding but is instead, a neural marker of endogenous processing related to abstract categories and learning (Alain *et al.*, 2007; Alain *et al.*, 2010; Bidelman *et al.*, 2013b; Ross *et al.*, 2013; Bidelman and Walker, 2017). The notion that smaller ERP responses correspond to better categorization for complex auditory stimuli has been reported in other training studies, and is often attributed to more efficient neural processing after short-term learning (Alain *et al.*, 2010; Ben-David *et al.*, 2011).

While these results show promise for understanding the psychobiological dynamics of auditory categorization, there are caveats and limitations of the current approach worth mentioning. Our models make several assumptions about the nature of learning and categorization based on prior work on CP (Harnad, 1987; Livingston *et al.*, 1998) and learner models in the GKT framework (Cen *et al.*, 2006; Pavlik Jr *et al.*, 2009; Chi *et al.*, 2011; Pavlik Jr *et al.*, 2020). For instance, we assume learning only occurs during overt training. Instead, implicit learning experiments suggest that categorization performance can improve through training without explicit knowledge of the categorical structure (Luthra *et al.*, 2019). However, our results indicate that categorization responses exhibit the greatest change during training where feedback on the interval category is present (Fig. 3). Second, the improvement in learning performance is captured by a logit parameter (“*Learn*”), the log of the ratio between a running sum of correct to incorrect trials (see Pavlik Jr *et al.*, 2020)^3^. This implies learning is perhaps related to monitoring ongoing successes and failures during the training paradigm. As such, these model assumptions may limit generalizability to other datasets, particularly those experiments without a training component.

Additionally, our models’ McFadden’s R^2^ values are smaller than other examples reported in the learner model literature (e.g., Pavlik Jr *et al.*, 2020), which indicates substantial variance in categorization performance not captured by even the best model configuration (i.e., > 85%). This could simply be the result of fitting a model to a very noisy behavioral task (i.e., lots of error). Alternatively, a larger dataset and a more complex task than our design might capture additional variance from other factors that contribute to identification performance. This could also allow for comparing individual student learning parameters without risk of overfitting as well as the inclusion of model parameters that, for example, capture initial, baseline knowledge about musical intervals prior to training (i.e., using separate intercepts for each learner) or different learning rates for each individual (i.e., using different learning slopes for each learner) (Cen *et al.*, 2006; Pavlik Jr *et al.*, 2009; Liu and Koedinger, 2017). Multiple days of training may also permit deeper probing of brain-behavior relationships supporting auditory category learning, including how different training regimens might enhance performance (Little *et al.*, 2019) or whether incorporating known psychological constructs like recency or forgetting/decay effects impact predicted learning outcomes (Pavlik Jr *et al.*, 2020).

Another important consideration of these models is that the outcome is a binary code for whether a subject responds minor (coded as 0) or major (coded as 1) rather than whether or not the subject was correct on a given trial, as is common in the learner modeling literature. This choice accommodated the subjectivity in auditory categorization; our task does not afford a true “correct” or “incorrect” response. While possible to reconfigure models for predicting “accuracy,” it would not be appropriate to consider identification of the inner tokens (particularly the ambiguous mid-continuum token) as “correct” or “incorrect” because these stimuli, by definition, do not fit neatly into a single category. Subjective categorization judgments also naturally differ across individuals, reminiscent of fuzzy logic models or gradient theories of categorization processes (e.g., Massaro, 1987; McMurray *et al.*, 2008). Only continuum endpoints (minor/major prototypes), could justifiably be used for determination of subject accuracy across the experiment, but this would miss the local transfer effects of categorizing inner 3^rds^ tokens not heard during training (and, apparently, their subsequent interference effects in the posttest; see Fig. 3). Our use of a continuum that is more graded rather than all-or-nothing (Medin and Barsalou, 1987) might limit generalizability to other studies that model category learning using stimuli with more binary, “true/false” category properties (e.g., visual shapes: Kruschke, 1992; bird species: Roads and Mozer, 2019).

Future studies might benefit from alternative methods of incorporating neural data in computational models to better understand categorical processes in auditory learning. For example, source localization techniques could be used to estimate the neural response from specific regions in the brain to identify the neural networks most important for learning and acquiring category structure (Liebenthal *et al.*, 2005; Desai *et al.*, 2008; Myers *et al.*, 2009; Chang *et al.*, 2010; Myers and Swan, 2012; Bidelman and Walker, 2019; Mankel *et al.*, 2020). Similarly, single-trial neural data could be used to understand mechanistic differences in speech vs. music categorization (Cutting and Rosner, 1974; Weidema *et al.*, 2016; Bidelman and Walker, 2017; 2019), long-term neuroplasticity (Siegel and Siegel, 1977; Burns and Ward, 1978; Zatorre and Halpern, 1979; Klein and Zatorre, 2011; Bidelman *et al.*, 2014; Bidelman and Alain, 2015; Bidelman and Lee, 2015; Wu *et al.*, 2015; Bidelman and Walker, 2019) attentional modulation (Bidelman and Walker, 2017), and individual differences in CP (Howard *et al.*, 1992; Mankel *et al.*, 2020).

## V. CONCLUSIONS

In conclusion, we demonstrate that computational learning models can be used to trace the rapid development of novel sound categories for musical intervals within ~20 minutes of feedback training. Moreover, neuroimaging data (single-trial EEG) helps decipher listeners’ behavioral gains in learning during training. Specifically, trial-by-trial changes in brain activity (P2 ~150 ms) were associated with greater probability of correct interval identification after learning. Our study highlights the more nuanced possibilities of adapting sophisticated computational models of learning theory to understand how dynamic coupling between brain and behavior drives the time course of auditory categorical learning.

## ACKNOWLEDGEMENTS

This work was supported by the National Institute on Deafness and Other Communication Disorders of the National Institutes of Health under award numbers NIH/NIDCD F31DC019041 (K.M.) and R01DC016267 (G.M.B.) as well as the National Science Foundation Data Infrastructure Building Blocks program under grant number ACI-1443068 (P.P., Jr.). Data are available from G.M.B. upon reasonable request.

---

**APPENDIX.**
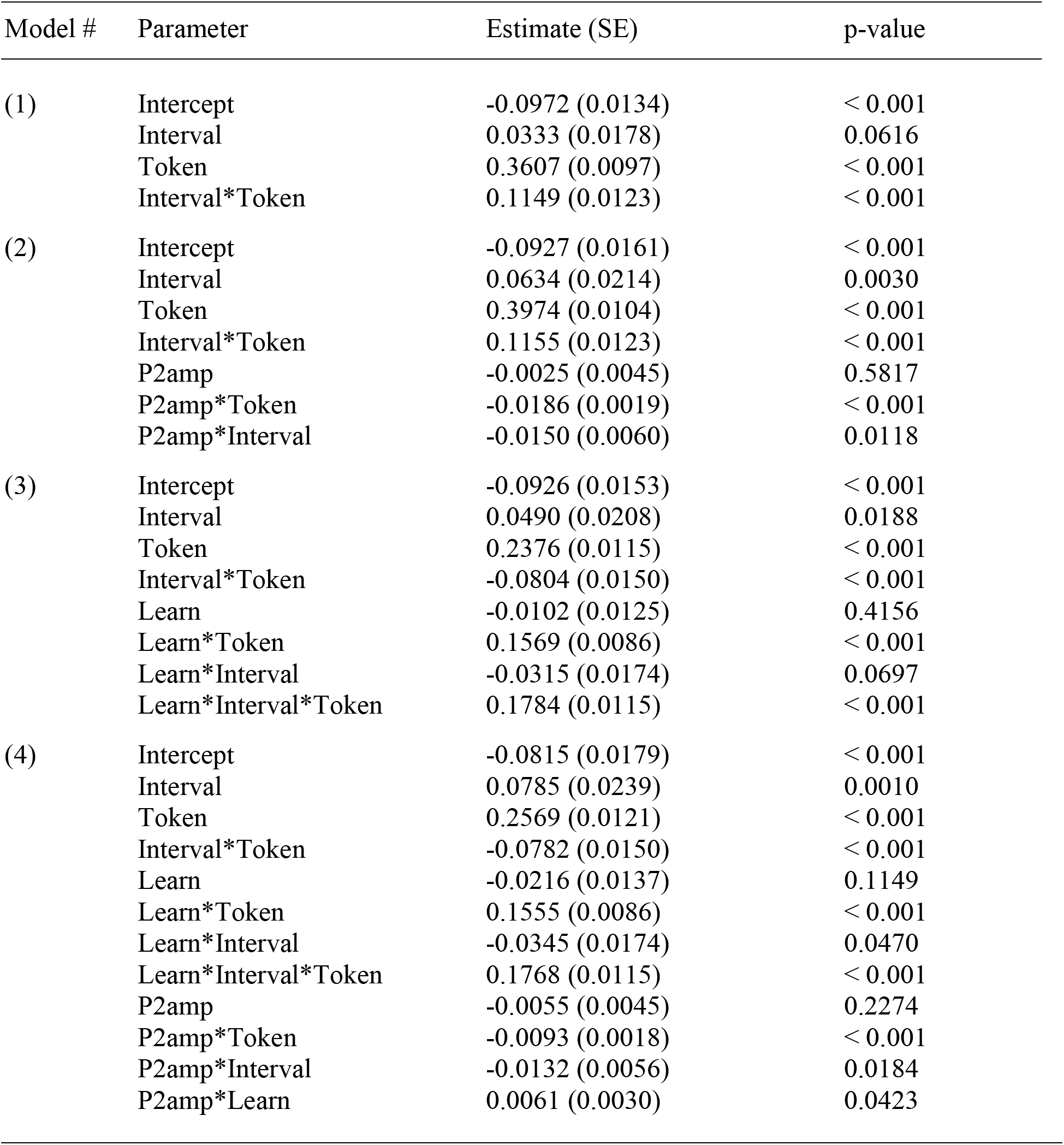
TABLE OF MODEL PARAMETER ESTIMATES

## TEXTUAL FOOTNOTES

1 One pilot subject heard 100 presentations of each token in the pre- and post-training phases. Due to technical issues, two participants terminated one of the blocks in the posttest early (though both were at least ~3/4 complete with the block), but since the missing data constituted <10% of their responses, imputation was not performed (Newman, 2014).

2 One pilot subject received only 6 blocks of training while another pilot subject was tested on 15 blocks of training before the final number of 10 blocks was settled for all others. All of these trials were included in the analyses. One subject’s EEG responses were not recorded during training, so their trials were excluded from learning model analyses.

3 During model building, a logit parameter offered a cleaner explanation of results by combining in a ratio the correct and incorrect trials rather than fitting separate parameters for each (Pavlik Jr *et al.*, 2009), and it provides diminishing marginal returns to learning for later vs. earlier training trials in accordance with theories of learning (Pavlik Jr *et al.*, 2020).

## References

Alain, C., Campeanu, S., and Tremblay, K. (2010). “Changes in sensory evoked responses coincide with rapid improvement in speech identification performance,” J. Cogn. Neurosci. 22, 392–403.

Alain, C., Snyder, J. S., He, Y., and Reinke, K. S. (2007). “Changes in auditory cortex parallel rapid perceptual learning,” Cereb. Cortex 17, 1074–1084.

Alho, J., Green, B. M., May, P. J. C., Sams, M., Tiitinen, H., Rauschecker, J. P., and Jääskeläinen, I. P. (2016). “Early-latency categorical speech sound representations in the left inferior frontal gyrus,” Neuroimage 129, 214–223.

Ben-David, B. M., Campeanu, S., Tremblay, K. L., and Alain, C. (2011). “Auditory evoked potentials dissociate rapid perceptual learning from task repetition without learning,” Psychophysiology 48, 797–807.

Bidelman, G. M., and Alain, C. (2015). “Musical training orchestrates coordinated neuroplasticity in auditory brainstem and cortex to counteract age-related declines in categorical vowel perception,” J. Neurosci. 35, 1240–1249.

Bidelman, G. M., Bush, L. C., and Boudreaux, A. M. (2020). “Effects of Noise on the Behavioral and Neural Categorization of Speech,” Front. Neurosci. 14, 153.

Bidelman, G. M., Hutka, S., and Moreno, S. (2013a). “Tone language speakers and musicians share enhanced perceptual and cognitive abilities for musical pitch: Evidence for bidirectionality between the domains of language and music,” PloS One 8, e60676.

Bidelman, G. M., and Lee, C.-C. (2015). “Effects of language experience and stimulus context on the neural organization and categorical perception of speech,” Neuroimage 120, 191–200.

Bidelman, G. M., Moreno, S., and Alain, C. (2013b). “Tracing the emergence of categorical speech perception in the human auditory system,” Neuroimage 79, 201–212.

Bidelman, G. M., and Walker, B. S. (2017). “Attentional modulation and domain-specificity underlying the neural organization of auditory categorical perception,” Eur. J. Neurosci. 45, 690–699.

Bidelman, G. M., and Walker, B. S. (2019). “Plasticity in auditory categorization is supported by differential engagement of the auditory-linguistic network,” Neuroimage 201, 1–10.

Bidelman, G. M., Weiss, M. W., Moreno, S., and Alain, C. (2014). “Coordinated plasticity in brainstem and auditory cortex contributes to enhanced categorical speech perception in musicians,” Eur. J. Neurosci. 40, 2662–2673.

Binder, J. R., Liebenthal, E., Possing, E. T., Medler, D. A., and Ward, B. D. (2004). “Neural correlates of sensory and decision processes in auditory object identification,” Nat. Neurosci. 7, 295–301.

Bouton, S., Chambon, V., Tyrand, R., Guggisberg, A. G., Seeck, M., Karkar, S., van de Ville, D., and Giraud, A. L. (2018). “Focal versus distributed temporal cortex activity for speech sound category assignment,” Proc. Natl. Acad. Sci. USA 115, E1299–E1308.

Burns, E. M., and Ward, W. D. (1978). “Categorical perception - phenomenon or epiphenomenon: Evidence from experiments in the perception of melodic musical intervals,” J. Acoust. Soc. Am. 63, 456–468.

Carcagno, S., and Plack, C. J. (2011). “Pitch discrimination learning: Specificity for pitch and harmonic resolvability, and electrophysiological correlates,” J. Assoc. Res. Oto. 12, 503–517.

Cen, H., Koedinger, K., and Junker, B. (2006). “Learning Factors Analysis: A General Method for Cognitive Model Evaluation and Improvement,” in International Conference on Intelligent Tutoring Systems, edited by M. Ikeda, K. D. Ashley, and T.-W. Chan (Springer, Jhongli, Taiwan), pp. 164–176.

Chang, E. F., Rieger, J. W., Johnson, K., Berger, M. S., Barbaro, N. M., and Knight, R. T. (2010). “Categorical speech representation in human superior temporal gyrus,” Nat. Neurosci. 13, 1428–1432.

Chi, M., Koedinger, K. R., Gordon, G., Jordan, P., and VanLehn, K. (2011). “Instructional Factors Analysis: A cognitive model for multiple instructional interventions,” in International Conference on Educational Data Mining (Eindhoven, Netherlands), pp. 61–70.

Cutting, J. E., and Rosner, B. S. (1974). “Categories and boundaries in speech and music,” Percept. Psychophys. 16, 564–570.

de Souza, A. C. S., Yehia, H. C., Sato, M.-a., and Callan, D. (2013). “Brain activity underlying auditory perceptual learning during short period training: simultaneous fMRI and EEG recording,” BMC Neurosci. 14, 8.

Desai, R., Liebenthal, E., Waldron, E., and Binder, J. R. (2008). “Left posterior temporal regions are sensitive to auditory categorization,” J. Cogn. Neurosci. 20, 1174–1188.

Draganova, R., Wollbrink, A., Schulz, M., Okamoto, H., and Pantev, C. (2009). “Modulation of auditory evoked responses to spectral and temporal changes by behavioral discrimination training,” BMC Neurosci. 10, 143.

Du, Y., He, Y., Arnott, S. R., Ross, B., Wu, X., Li, L., and Alain, C. (2015). “Rapid tuning of auditory “what” and “where” pathways by training,” Cereb. Cortex 25, 496–506.

Galyardt, A., and Goldin, I. (2015). “Move your lamp post: Recent data reflects learner knowledge better than older data,” Journal of Educational Data Mining 7, 83–108.

Golestani, N., and Zatorre, R. J. (2004). “Learning new sounds of speech: Reallocation of neural substrates,” Neuroimage 21, 494–506.

Gong, Y., Beck, J. E., and Hefferman, N. T. (2011). “How to construct more accurate student models: Comparing and optimizing knowledge tracing and performance factor analysis,” International Journal of Artificial Intelligence in Education 21, 27–46.

Guenther, F. H., Husain, F. T., Cohen, M. A., and Shinn-Cunningham, B. G. (1999). “Effects of categorization and discrimination training on auditory perceptual space,” J. Acoust. Soc. Am. 106, 2900–2912.

Guenther, F. H., Nieto-Castanon, A., Ghosh, S. S., and Tourville, J. A. (2004). “Representation of sound categories in auditory cortical maps,” J. Speech. Lang. Hear. Res. 47, 46–57.

Hall, J. W. (1992). Handbook of Auditory Evoked Responses (Allyn and Bacon, Needham Heights).

Harnad, S. (1987). Categorical perception: The groundwork of cognition (Cambridge University Press, New York).

Howard, D., Rosen, S., and Broad, V. (1992). “Major/Minor triad identification and discrimination by musically trained and untrained listeners,” Music Percept. 10, 205–220.

Khajah, M. M., Lindsey, R. V., and Mozer, M. C. (2014). “Maximizing Students’ Retention via Spaced Review: Practical Guidance From Computational Models of Memory,” Topics in Cognitive Science 6, 157–169.

Klein, M. E., and Zatorre, R. J. (2011). “A role for the right superior temporal sulcus in categorical perception of musical chords,” Neuropsychologia 49, 878–887.

Klein, M. E., and Zatorre, R. J. (2015). “Representations of invariant musical categories are decodable by pattern analysis of locally distributed BOLD responses in superior temporal and intraparietal sulci,” Cereb. Cortex 25, 1947–1957.

Kraus, N., McGee, T., Carrell, T., King, C., Tremblay, K., and Nicol, T. (1995). “Central auditory system plasticity associated with speech discrimination training,” J. Cogn. Neurosci. 7, 25–32.

Kruschke, J. K. (1992). “ALCOVE: An exemplar-based connectionist model of category learning,” Psychol. Rev. 99, 22–44.

Kuhl, P. K. (1991). “Human adults and human infants show a “perceptual magnet effect” for the prototypes of speech categories, monkeys do not,” Percept. Psychophys. 50, 93–107.

Kuhl, P. K., Williams, K. A., Lacerda, F., Stevens, K. N., and Lindblom, B. (1992). “Linguistic experience alters phonetic perception in infants by 6 months of age,” Science 255, 606–608.

Lee, R. R. W., Hsu, C. H., Lin, S. K., Wu, D. H., and Tzeng, O. J. L. (2017). “Learning transforms functional organization for Mandarin lexical tone discrimination in the brain: Evidence from a MEG experiment on second language learning,” J. Neuroling. 42, 124–139.

Lee, Y.-S., Turkeltaub, P., Granger, R., and Raizada, R. D. S. (2012). “Categorical speech processing in Broca’s area: An fMRI study using multivariate pattern-based analysis,” J. Neurosci. 32, 3942–3948.

Li, P., Sepanski, S., and Zhao, X. (2006). “Language history questionnaire: A Web-based interface for bilingual research,” Behav. Res. Meth. 38, 202–210.

Liebenthal, E., Binder, J. R., Spitzer, S. M., Possing, E. T., and Medler, D. A. (2005). “Neural substrates of phonemic perception,” Cereb Cortex 15, 1621–1631.

Liebenthal, E., Desai, R., Ellingson, M. M., Ramachandran, B., Desai, A., and Binder, J. R. (2010). “Specialization along the left superior temporal sulcus for auditory categorization,” Cereb Cortex 20, 2958–2970.

Little, D. F., Cheng, H. H., and Wright, B. A. (2019). “Inducing musical-interval learning by combining task practice with periods of stimulus exposure alone,” Atten Percept Psychophys 81, 344–357.

Liu, R., and Koedinger, K. R. (2017). “Towards reliable and valid measurement of individualized student parameters,” in International Conference on Educational Data Mining, edited by X. Hu, T. Barnes, A. Hershkovitz, and L. Paquette (Wuhan, China), pp. 135–142.

Lively, S. E., Logan, J. S., and Pisoni, D. B. (1993). “Training Japanese listeners to identify English /r/ and /l/: II. The role of phonetic environment and talker variability in learning new perceptual categories,” J. Acoust. Soc. Am. 94, 1242–1255.

Livingston, K. R., Andrews, J. K., and Harnad, S. (1998). “Categorical perception effects induced by category learning,” J. Exp. Psychol.-Learn. Mem. Cogn. 24, 732–753.

Luck, S. (2014). “The design of ERP experiments,” in An introduction to the event-related potential technique (MIT Press, Cambridge, MA), pp. 119–146.

Luthra, S., Fuhrmeister, P., Molfese, P. J., Guediche, S., Blumstein, S. E., and Myers, E. B. (2019). “Brain-behavior relationships in incidental learning of non-native phonetic categories,” Brain Lang. 198, 104692.

Mankel, K., Barber, J., and Bidelman, G. M. (2020). “Auditory categorical processing for speech is modulated by inherent musical listening skills,” NeuroReport 31, 162–166.

Massaro, D. M. (1987). “Categorical partition: A fuzzy-logical model of categorization behavior,” in Categorical perception: The groundwork of cognition, edited by S. Harnad (Cambridge University Press, New York), pp. 254–283.

McMurray, B., Aslin, R. N., Tanenhaus, M. K., Spivey, M. J., and Subik, D. (2008). “Gradient sensitivity to within-category variation in words and syllables,” J. Exp. Psychol. Hum. Percept. Perform. 34, 1609–1631.

Medin, D. L., and Barsalou, L. W. (1987). “Categorization processes and categorical perception,” in Categorical perception: The groundwork of cognition, edited by S. Harnad (Cambridge University Press, New York), pp. 455–490.

Myers, E. B., Blumstein, S. E., Walsh, E., and Eliassen, J. (2009). “Inferior frontal regions underlie the perception of phonetic category invariance,” Psych. Sci. 20, 895–903.

Myers, E. B., and Swan, K. (2012). “Effects of category learning on neural sensitivity to non-native phonetic categories,” J. Cogn. Neurosci. 24, 1695–1708.

Newman, D. A. (2014). “Missing Data: Five Practical Guidelines,” Organizational Research Methods 17, 372–411.

Oldfield, R. C. (1971). “The assessment and analysis of handedness: The Edinburgh inventory,” Neuropsychologia 9, 97–113.

Oostenveld, R., and Praamstra, P. (2001). “The five percent electrode system for high-resolution EEG and ERP measurements,” Clin. Neurophysiol. 112, 713–719.

Pavlik Jr, P. I., Cen, H., and Koedinger, K. R. (2009). “Performance factors analysis: A new alternative to knowledge tracing,” in 14th International Conference on Artificial Intelligence in Education, edited by V. Dimitrova, R. Mizoguchi, B. D. Boulay, and A. Graesser (Brighton, England).

Pavlik Jr, P. I., Eglington, L. G., and Harrell-Williams, L. M. (2020). “Generalized Knowledge Tracing: A constrained framework for learner modeling,” arXiv (stat.AP).

Pavlik Jr, P. I., Hua, H., Williams, J., and Bidelman, G. M. (2013). “Modeling the effect of spacing on musical interval training,” in Proceedings of 6th International Conference on Educational Data Mining (July 6-9, Memphis, TN).

Pavlik Jr., P. I., and Anderson, J. R. (2008). “Using a model to compute the optimal schedule of practice,” Journal of Experimental Psychology: Applied 14, 101–117.

Pisoni, D. B., Aslin, R. N., Perey, A. J., and Hennessy, B. L. (1982). “Some effects of laboratory training on identification and discrimination of voicing contrasts in stop consonants,” J. Exp. Psychol.-Hum. Percept. Perform. 8, 297–314.

Roads, B. D., and Mozer, M. C. (2019). “Predicting the difficulty of human category learning using exemplar-based neural networks,” Manuscript submitted for publication.

Rosen, S., and Howell, P. (1987). “Auditory, articulatory and learning explanations of categorical perception in speech,” in Categorical perception: The groundwork of cognition, edited by S. R. Harnad (Cambridge University Press, New York), pp. 113–160.

Ross, B., Jamali, S., and Tremblay, K. L. (2013). “Plasticity in neuromagnetic cortical responses suggests enhanced auditory object representation,” BMC Neurosci. 14, 151.

Siegel, J. A., and Siegel, W. (1977). “Categorical perception of tonal intervals - musicians can’t tell sharp from flat,” Percept. Psychophys. 21, 399–407.

Spada, H., and McGaw, B. (1985). “The assessment of learning effects with linear logistic test models,” in Developments in psychology and psychometrics, edited by S. Embretson (Academic Press, Orlando, FL).

Swan, K., and Myers, E. (2013). “Category labels induce boundary-dependent perceptual warping in learned speech categories,” Second Lang. Res. 29, 391–411.

Tong, Y., Melara, R. D., and Rao, A. (2009). “P2 enhancement from auditory discrimination training is associated with improved reaction times,” Brain Res. 1297, 80–88.

Toscano, J. C., Anderson, N. D., Fabiani, M., Gratton, G., and Garnsey, S. M. (2018). “The time-course of cortical responses to speech revealed by fast optical imaging,” Brain Lang. 184, 32–42.

Tremblay, K., Kraus, N., McGee, T., Ponton, C., and Otis, B. (2001). “Central auditory plasticity: Changes in the N1-P2 complex after speech-sound training,” Ear Hear. 22, 79–90.

Weidema, J. L., Roncaglia-Denissen, M. P., and Honing, H. (2016). “Top-down modulation on the perception and categorization of identical pitch contours in speech and music,” Front. Psychol. 7, 817.

Wu, H., Ma, X., Zhang, L., Liu, Y., Zhang, Y., and Shu, H. (2015). “Musical experience modulates categorical perception of lexical tones in native Chinese speakers,” Front. Psychol. 6, 436.

Yarkoni, T., and Westfall, J. (2017). “Choosing Prediction Over Explanation in Psychology: Lessons From Machine Learning,” Perspectives on psychological science: a journal of the Association for Psychological Science 12, 1100–1122.

Yi, H. G., and Chandrasekaran, B. (2016). “Auditory categories with separable decision boundaries are learned faster with full feedback than with minimal feedback,” Journal of the Acoustic Society of America 140, 1332.

Zatorre, R. J., and Halpern, A. R. (1979). “Identification, discrimination, and selective adaptation of simultaneous musical intervals,” Percept. Psychophys. 26, 384–395.

